# Association between mitochondrial DNA copy number and production traits in pigs

**DOI:** 10.1101/2024.02.07.579287

**Authors:** Eduard Molinero, Ramona N. Pena, Joan Estany, Roger Ros-Freixedes

## Abstract

Mitochondria are essential organelles in the regulation of cellular energetic metabolism. Mitochondrial DNA copy number (mtDNA_CN) can be used as a proxy for mitochondria number, size, and activity. The aims of our study are to evaluate the effect of mtDNA_CN and mitochondrial haploblocks on production traits in pigs, and to identify the genetic background of this cellular phenotype. We collected performance data of 234 pigs and extracted DNA from skeletal muscle. Whole-genome sequencing data was used to determine mtDNA_CN. We detected positive correlations of muscle mtDNA_CN with backfat thickness at 207 d (+0.14; *p*-value = 0.07), and negative correlations with carcase loin thickness (–0.14; *p*-value = 0.03). Pigs with less mtDNA_CN had greater loin thickness (+4.1 mm; *p*-value = 0.01) and lower backfat thickness (–1.1 mm; *p*-value = 0.08), which resulted in greater carcase lean percentage (+2.4%; *p*-value = 0.04) than pigs with high mtDNA_CN. These results support the hypothesis that a reduction of mitochondrial activity is associated with greater feed efficiency. Pork from pigs with less mtDNA_CN may also tend to have lower ultimate pH (correlation: +0.19; *p*-value < 0.01). We found no association of the most frequent mitochondrial haploblocks with mtDNA_CN or the production traits, but several genomic regions that harbour potential candidate genes with functions related to mitochondrial biogenesis and homeostasis were associated to mtDNA_CN. These regions provide new insights into the genetic background of this cellular phenotype but it is still uncertain if such associations translate into noticeable effects on the production traits.

## Introduction

Mitochondria are subcellular organelles whose main function is to generate energy for the cells, as well as being involved in many other metabolic pathways, which include signalling, ion homeostasis, lipid metabolism, redox control, cell growth, and cell death (McBride, Neuspiel and Wasiak, 2006). Mitochondria provide 90% of the energy needed for cell metabolism (Emmerson, 1997), generated in their membrane by the electron transport chain proteins through oxidative phosphorylation of adenosine diphosphate (ADP) into adenosine triphosphate (ATP) (McBride, Neuspiel and Wasiak, 2006). It has been hypothesised that defective proteins in the electron transport chain could lead to a reduction of energy efficiency of animals (Bottje, 2019). Suboptimal mitochondrial function could lead to an increase of reactive oxygen species (Tait and Green, 2012), which might start a cascade of oxidative damage to proteins, and specifically to the electron transporter chain, further reducing energy production (Iqbal *et al*., 2005).

Several studies reported a relationship between mitochondrial activity and feed efficiency in the context of livestock production. Bottje *et al*. (2002) reported that less feed-efficient broilers had increased electron leak in breast mitochondria and lower activities of protein complexes I and II. This study was followed by several more in poultry (Iqbal *et al*., 2005; Lassiter *et al*., 2006; Kong *et al*., 2016; Bottje *et al*., 2017), cattle (Kelly *et al*., 2011; Dorji *et al*., 2021), and pigs (Grubbs *et al*., 2013; Jing *et al*., 2015; Fu *et al*., 2017; Gondret *et al*., 2017; Carmelo and Kadarmideen, 2020). Most studies have concluded that individuals with high feed efficiency show a reduction of the expression of mitochondrial genes (Iqbal *et al*., 2005; Dorji *et al*., 2021), as well as a reduction of the expression of genes involved in mitochondrial biogenesis (Jing *et al*., 2015) and in oxidative phosphorylation (Fu *et al*., 2017). Nevertheless, there is controversy about these results, as other researchers have reported contradictory results, such as the increase of expression (Grubbs *et al*., 2013; Gondret *et al*., 2017) and activity (Bottje *et al*., 2002) of electron transport chain proteins. Thus, the direction of the association between mitochondrial activity and feed efficiency remains inconclusive and seems dependent on the species and sampled tissue.

Mitochondrial activity can be measured using laboratory methodologies such as the determination of oxygen consumption rate or the production of ATP or reactive oxygen species. However, mitochondrial DNA (mtDNA) copy number (mtDNA_CN), which can be easily derived from raw whole-genome sequence data, can be used as a proxy for mitochondrial activity. D’Erchia *et al*. (2015) found that mtDNA_CN was positively correlated with mitochondrial mass and the respiratory capacity. Variation in mtDNA_CN has been associated with diverse pathologies, such as cardiomyopathy, diabetes and cancer (Kaaman *et al*., 2007; Dillon *et al*., 2012; Holmström *et al*., 2012; Pirinen *et al*., 2020; Abd Radzak *et al*., 2022). This cellular trait is moderately heritable, with heritability estimates in humans of 0.33 to 0.65 (Xing *et al*., 2008; López *et al*., 2012). Two studies in livestock species have associated mtDNA_CN with carcase and breast meat yield and abdominal fat, in chickens (Reverter *et al*., 2016), and birth and weaning weight, in cattle (Sanglard *et al*., 2023). To our knowledge, there is no information about the relationship between mtDNA_CN and production traits in pigs.

The aims of this study were: (i) to evaluate the effect of mtDNA_CN on growth, carcase composition, and meat quality traits in pigs, (ii) to evaluate the effect of mitochondrial haploblocks on the same traits, and (iii) to identify the genetic background of this cellular phenotype.

## Materials and Methods

### Animals and phenotypes

A total of 234 Duroc pigs from the same genetic line were raised in 19 batches between 2002 and 2021 following a common protocol for data recording and tissue sampling (Ros-Freixedes *et al*., 2016). Pigs from each batch were raised from around 75 days until slaughter age (at 213 days, SD 7.8 d) under identical conditions. During this time, animals had ad libitum access to commercial feed (Esporc, Riudarenes, Spain). At 207 days of age (SD 8 d), pigs were weighed and the backfat and loin thickness were measured at 5 cm off the midline at the position of the last rib using a portable ultrasonic scanner (Piglog 105; Frontmatec, Kolding, Denmark). All pigs were slaughtered in the same abattoir, where carcase weight was recorded. Backfat and loin thickness at 6 cm off the midline between the third and fourth last ribs were recorded using an on-line ultrasound automatic scanner (AutoFOM; Frontmatec, Kolding, Denmark). Ultimate pH of meat was measured in muscle *semimembranosus* after chilling for approximately 24 h at 2°C with a pH-meter equipped with a spear-tipped probe (Testo 205, Testo AG, Lenzkirch, Germany). Then, for a subset of 205 pigs (15 batches), samples of muscle *gluteus medius* were taken, vacuum packaged and stored at –20 °C until required for chemical analyses. Intramuscular fat content and fatty acid composition was determined in duplicate by quantitative gas chromatography (Bosch *et al*., 2009). Fatty acids were expressed as percentages relative to total fatty acid content. The proportion of saturated fatty acids (C14:0, C16:0, C18:0 and C20:0), monounsaturated fatty acids (C16:1n-7, C18:1n-7, C18:1n-9 and C20:1n-9), and polyunsaturated fatty acids (C18:2n-6, C18:3n-3, C20:2n-6 and C20:4n-6) were calculated.

### Whole-genome sequencing

Extraction of DNA from all pigs was carried out following a conventional phenol:chloroform protocol (Green and Sambrook, 2017) from skeletal muscle (n=234). An additional set of 48 individuals, from the last 5 batches (between 2019 and 2021) and that followed the same rearing and sampling protocol as the other 234 pigs, had DNA extracted from blood at 85.6 days of age (SD 2.4 d) to assess the value of non-target tissues for mtDNA_CN determination.

The DNA samples were submitted to the Centre Nacional d’Anàlisi Genòmica (CNAG-CRG, Barcelona, Spain) for whole-genome sequencing on a NovaSeq 6000 instrument (Illumina, San Diego, CA, USA) in paired-end mode, as described by Molinero *et al*. (2022).

Trimmomatic (Bolger, Lohse and Usadel, 2014) was used to remove adapter sequences from the FASTQ sequence files. The trimmed reads were aligned to the reference genome *Sscrofa11.1* (GCA_000003025.6; Warr *et al*., 2020) using the BWA-MEM algorithm (Li, 2013). The average realized sequencing coverage depth was 7.3x (SD 2.0x). After using Picard (http://broadinstitute.github.io/picard) to tag duplicates, the variant calling was performed separately for each individual with GATK HaplotypeCaller (GATK 3.8.0) (DePristo *et al*., 2011) using default settings. Then, all individuals were jointly genotyped for all variant positions. We retained all biallelic variants, including single nucleotide polymorphisms and short insertions/deletions, for further analyses with VCFtools (Danecek *et al*., 2011).

The mtDNA_CN was calculated as the natural logarithm of the ratio between mitochondrial and nuclear DNA coverage depths. Mitochondrial haploblocks were obtained using Haploview 4.2 (Barrett *et al*., 2005) with the standard method described by Gabriel *et al*. (2002).

### Relationship with production traits and meat quality

We used a linear model to adjust mtDNA_CN and each trait phenotypes for the batch and covariate (if applicable) effects. For carcase weight we included the age of slaughter as a covariate, while for the other production traits we included carcase weight as a covariate. For fatty acid composition traits, carcase weight and intramuscular fat content were both included as covariates. Then, we tested the phenotypic correlations between the preadjusted values of mtDNA_CN and the different production traits. We also generated two divergent groups for mtDNA_CN by selecting the 50 animals with the highest and 50 with the lowest values amongst the 205 pigs with complete fat content and fatty acid composition data. These two groups were subsequently used to study potential associations between mtDNA_CN and production traits. The difference between the two mtDNA_CN groups for each trait was tested with an *F*-test using a linear model that also included the effects of batch and covariates as described above. The effect of mitochondrial haploblocks and the rs709596309 leptin receptor (*LEPR*) genotypes (Ros-Freixedes et la., 2016) were similarly tested with models that included the same effects. All analyses were performed using the statistical package JMP PRO 16 (SAS Institute Inc., Cary, NC).

### Genome-wide association study

A genome-wide association study (GWAS) for mtDNA_CN, as a cellular phenotype using data from all 234 pigs sequenced from skeletal muscle, was performed using a univariate linear mixed model that accounted for the genomic relationship matrix. The following model was fitted using GEMMA 0.96 software (Zhou and Stephens, 2012) to evaluate the association between the phenotypes and each SNP:

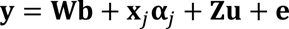

where **y** is the vector of phenotypic observations; **W** is the incidence matrix for fixed effects; **b** is the vector of the fixed effects, including an intercept and the batch effect; 𝐱_𝑗_is the vector of genotypes of the j^th^ variant coded as 0 and 2 for homozygotes and 1 for heterozygotes; 𝛂_𝑗_ is the allele substitution effect of the j^th^ variant; 𝐙 is the incidence matrix for the random individual polygenic effects; 𝐙 is the vector of the random individual polygenic effects; and 𝐞𝐞 is the vector of residual terms. The random individual polygenic effects and residual terms were assumed to follow normal distributions 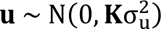 and 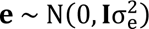, respectively, where 𝐊 is the genomic relationship matrix, 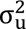 is the additive genetic variance, 𝐈 is an identity matrix, and 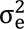 is the residual variance. Variants that had a *p*-value < 10^-5^ were taken as associated, and regions of 50 kb up- and downstream of the associated variants were explored for candidate genes. Overlapping regions arising from nearby significant variants were merged into a single wider region. Biomart tool in Ensembl (Martin *et al*., 2023) was used to retrieve candidate genes and their functions were annotated.

## Results

### Association between mtDNA_CN and production traits

Phenotypic correlations between mtDNA_CN and production traits are shown in Table 1. We detected positive correlations of muscle mtDNA_CN with pH (+0.19; *p*-value < 0.01) and backfat thickness at 207 d (+0.14; *p*-value = 0.07), and negative correlations with carcase loin thickness (–0.14; *p*-value = 0.03). Phenotypic correlations with fatty acid composition in muscle *gluteus medius* are shown in Table 2. We observed no correlation of mtDNA_CN with any of those traits.

**Table 1.** Phenotypic correlations of mtDNA_CN with production traits by tissue of DNA origin.

| Trait | Muscle<br>(n=234) | *p*-value | Blood<br>(n=48) | *p*-value |
| --- | --- | --- | --- | --- |
| Live measurements |  |  |  |  |
| Body weight<sup>a</sup> | −0.09 | 0.23 | +0.09 | 0.56 |
| Backfat thickness at 207 d | +0.14 | **0.07** | — | — |
| Loin thickness at 207 d | −0.06 | 0.48 | — | — |
| Carcase measurements |  |  |  |  |
| Carcase weight | −0.06 | 0.37 | +0.16 | 0.31 |
| Backfat thickness | +0.08 | 0.26 | +0.10 | 0.53 |
| Loin thickness | −0.14 | **0.03** | −0.06 | 0.69 |
| Carcase lean percentage | −0.09 | 0.19 | −0.05 | 0.74 |
| Intramuscular fat content | −0.02 | 0.76 | — | — |
| pH | +0.19 | **<0.01** | +0.13 | 0.43 |

<sup>a</sup> Body weight was registered at 207 and 85 days for muscle and blood, respectively.
In bold, *p*-values < 0.10.

**Table 2.** Phenotypic correlations of mtDNA_CN with fatty acid composition of muscle *gluteus medius* (% of total fatty acids).

| Trait | Correlation | *p*-value |
| --- | --- | --- |
| Saturated fatty acids | −0.01 | 0.95 |
| C14:0 | −0.05 | 0.52 |
| C16:0 | +0.03 | 0.60 |
| C18:0 | −0.03 | 0.63 |
| Monounsaturated fatty acids | +0.06 | 0.36 |
| C16:1n-7 | +0.05 | 0.45 |
| C18:1n-7 | +0.13 | 0.12 |
| C18:1n-9 | +0.04 | 0.61 |
| Polyunsaturated fatty acids | −0.09 | 0.19 |
| C18:2n-6 | −0.08 | 0.29 |
| C18:3n-3 | −0.08 | 0.24 |
| C20:4n.6 | −0.09 | 0.18 |

In bold, *p*-values < 0.10.

In order to better quantify the effect of mtDNA_CN on the production traits, we compared two groups of animals with the highest and lowest mtDNA_CN scores. The High group had an average mtDNA_CN score of 6.21 (SD 0.15) and the Low group had an average of 5.25 (SD 0.17). In line with the correlation data, we observed differences between the two groups for carcase composition traits (Table 3). The pigs with low mtDNA_CN had greater loin thickness (+4.1 mm; *p*-value = 0.01) and lower backfat thickness (–1.1 mm; *p*-value = 0.08), which resulted in greater carcase lean percentage (+2.4%; *p*-value = 0.04) than pigs with high mtDNA_CN. Similar trends were observed in the live body composition traits, although not significant. Inconsistencies between correlations for live body composition traits and carcase composition traits may derive from calibration biases of the instruments that were used to measure these traits at each stage, particularly when applied to pigs with high body fat content. However, both point towards pigs with more mtDNA_CN being leaner. No association between mtDNA_CN groups was detected for carcase weight, pH, or intramuscular fat content and fatty acid composition traits, except for a suggestive trend for the pigs with low mtDNA_CN to contain more polyunsaturated fat (Table 4).

**Table 3.** Least-square means (± standard errors) for production traits by mtDNA_CN.

| Trait | Low | High | *p*-value |
| --- | --- | --- | --- |
| Live measurements at 207 d |  |  |  |
| Body weight, kg | 127.3±2.0 | 125.2±2.1 | 0.49 |
| Backfat thickness, mm | 21.4±0.6 | 22.7±0.7 | 0.19 |
| Loin thickness, mm | 48.9±0.8 | 48.3±1.0 | 0.71 |
| Carcase measurements |  |  |  |
| Carcase weight, kg | 98.1±1.3 | 97.3±1.5 | 0.69 |
| Backfat thickness, mm | 22.6±0.4 | 23.7±0.5 | **0.08** |
| Loin thickness, mm | 47.6±1.1 | 43.5±1.2 | **0.01** |
| Carcase lean percentage, % | 43.1±0.8 | 40.7±0.9 | **0.04** |
| Intramuscular fat content, % | 17.8±0.8 | 18.6±0.9 | 0.42 |
| pH | 5.77±0.04 | 5.81±0.04 | 0.49 |

In bold, *p*-values < 0.10.

**Table 4.** Least-square means (± standard errors) for fatty acid composition (% of total fatty acids) in muscle *gluteus medius* by mtDNA_CN.

| Trait | Low | High | *p*-value |
| --- | --- | --- | --- |
| Saturated fatty acids | 37.9±0.3 | 38.1±0.3 | 0.63 |
| C14:0 | 1.74±0.05 | 1.68±0.05 | 0.37 |
| C16:0 | 24.3±0.2 | 24.5±0.2 | 0.58 |
| C18:0 | 11.7±0.2 | 11.8±0.2 | 0.54 |
| Monounsaturated fatty acids | 49.3±0.3 | 49.6±0.3 | 0.52 |
| C16:1n-7 | 3.56±0.09 | 3.57±0.09 | 0.97 |
| C18:1n-7 | 4.45±0.06 | 4.52±0.07 | 0.47 |
| C18:1n-9 | 41.4±0.3 | 41.5±0.3 | 0.67 |
| Polyunsaturated fatty acids | 12.8±0.2 | 12.3±0.2 | **0.06** |
| C18:2n-6 | 10.2±0.2 | 9.9±0.2 | 0.15 |
| C18:3n-3 | 0.59±0.01 | 0.55±0.02 | **0.07** |
| C20:4n.6 | 1.40±0.05 | 1.28±0.05 | 0.12 |

In bold, *p*-values < 0.10.

We previously described that the recessive missense variant rs709596309, which is a missense variant of the *LEPR* gene, is associated with feed intake, mobilization of energy reserves, and fat deposition in this population (Ros-Freixedes *et al*., 2016; Solé *et al*., 2021). We confirmed that the *LEPR* genotype affected carcase backfat thickness, loin thickness, and lean percentage in the sequenced pigs, but we did not find any evidence that these effects are underlain by any differences in mtDNA_CN (Table 5).

**Table 5.** Least-square means (± standard errors) for productions traits by *LEPR* genotypes.

| Trait | C– | TT | *p*-value |
| --- | --- | --- | --- |
| mtDNA_CN | 5.78±0.03 | 5.77±0.04 | 0.92 |
| Carcase measurements |  |  |  |
| Carcase weight, kg | 99.1±0.8 | 99.4±1.1 | 0.79 |
| Backfat thickness, mm | 24.0±0.3 | 25.9±0.4 | **<0.01** |
| Loin thickness, mm | 44.6±0.7 | 40.7±1.0 | **<0.01** |
| Carcase lean percentage, % | 42.3±0.5 | 37.9±0.7 | **<0.01** |
| Intramuscular fat content, % | 18.1±0.4 | 19.5±0.6 | **0.05** |
| pH | 5.77±0.02 | 5.80±0.03 | 0.39 |

In bold, *p*-values < 0.10.

The tissue that was used for extracting DNA had a clear effect on the estimated mtDNA_CN. Using raw sequencing data from DNA extracted from skeletal muscle resulted in greater mtDNA_CN (5.79; SD 0.37) than from blood samples (4.09; SD 0.34; *p*-value < 0.01). However, we observed similar correlation structures for mtDNA_CN estimated from skeletal muscle or blood samples, at least for pH and carcase composition traits. Due to the small sample size, the correlations for blood samples were not significant, yet the sign of the correlation for these traits was largely maintained with respect to skeletal muscle (Table 1).

### Effect of mtDNA haploblocks on traits

Twenty-six mitochondrial variants, all SNPs, segregated across the sequenced pigs. These variants formed 7 mitochondrial haploblocks, of which 4 had a frequency equal or greater than 0.10 and represented 98% of our population (Tables 6 and S1). No association was detected between the common haploblocks and mtDNA_CN or the production traits (Table 6).

**Table 6.** Least-square means (± standard errors) for production traits by mitochondrial haploblocks.

| Trait | Hap1 | Hap2 | Hap3 | Hap4 | *p*-value |
| --- | --- | --- | --- | --- | --- |
| Frequency | 0.41 | 0.30 | 0.17 | 0.10 | – |
| mtDNA_CN | 5.71±0.04 | 5.69±0.05 | 5.75±0.06 | 5.67±0.08 | 0.81 |
| Carcase measurements |  |  |  |  |  |
| Carcase weight, kg | 98.6±1.0 | 100.3±1.2 | 99.1±1.5 | 98.7±1.9 | 0.72 |
| Backfat thickness, mm | 23.3±0.3 | 23.6±0.4 | 24.3±0.5 | 23.4±0.6 | 0.37 |
| Loin thickness, mm | 43.9±0.9 | 44.7±1.1 | 44.1±1.4 | 43.8±1.8 | 0.94 |
| Carcase lean percentage, % | 42.0±0.7 | 41.5±0.8 | 39.7±1.0 | 41.6±1.3 | 0.28 |
| Intramuscular fat content, % | 19.0±0.5 | 18.5±0.6 | 18.1±0.8 | 18.3±1.0 | 0.77 |
| pH | 5.79±0.02 | 5.79±0.03 | 5.80±0.04 | 5.74±0.05 | 0.73 |

### Genome-wide association study for mtDNA_CN

Over 20 million variants were called in the Duroc pigs. We discarded variants that presented a genotyping rate below 0.9 and a minor allele frequency below 0.2 so that all genotypes were sufficiently represented in the 234 pigs. After the filtering, 7.05 million variants remained. The GWAS for mtDNA_CN identified 27 variants at *Sus scrofa* chromosomes (SSC) 3, 4, 5, 7, 10, 11, and 13 (Figure 1). Eleven of them were located at genomic regions where no annotated genes have been described. The remaining 16 were grouped into 9 genomic regions (Table 7) at SSC 3 (3 regions), 4, 5, and 10 (4 regions). A literature search pinpointed several candidate genes that have been previously associated with mitochondria homeostasis and activity, such as *ALODA*, *MAPK3*, *MTHFD2*, *HFM1*, several genes of the keratin (*KRT*) gene family, *HLX*, and *NEBL*.

**Figure 1.**
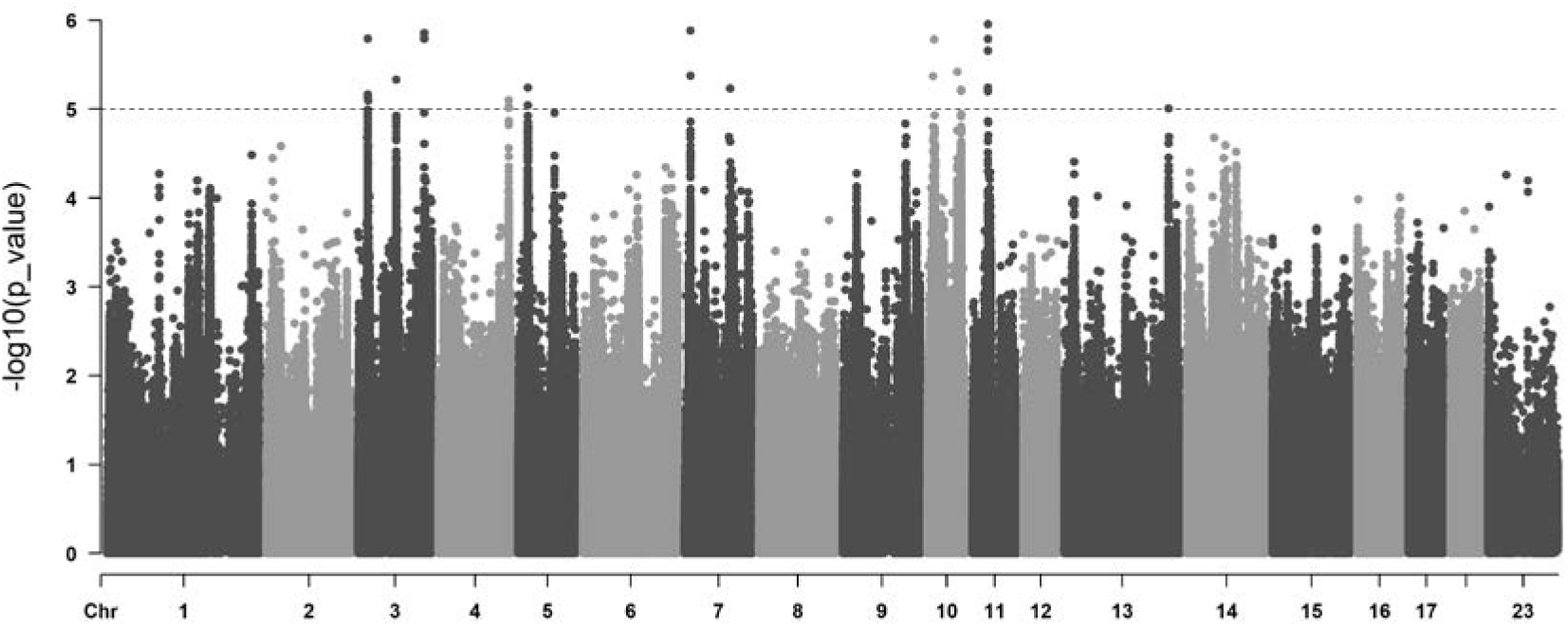
Manhattan plot using next generation sequencing data of 234 Duroc pigs for mtDNA_CN.

**Table 7.**
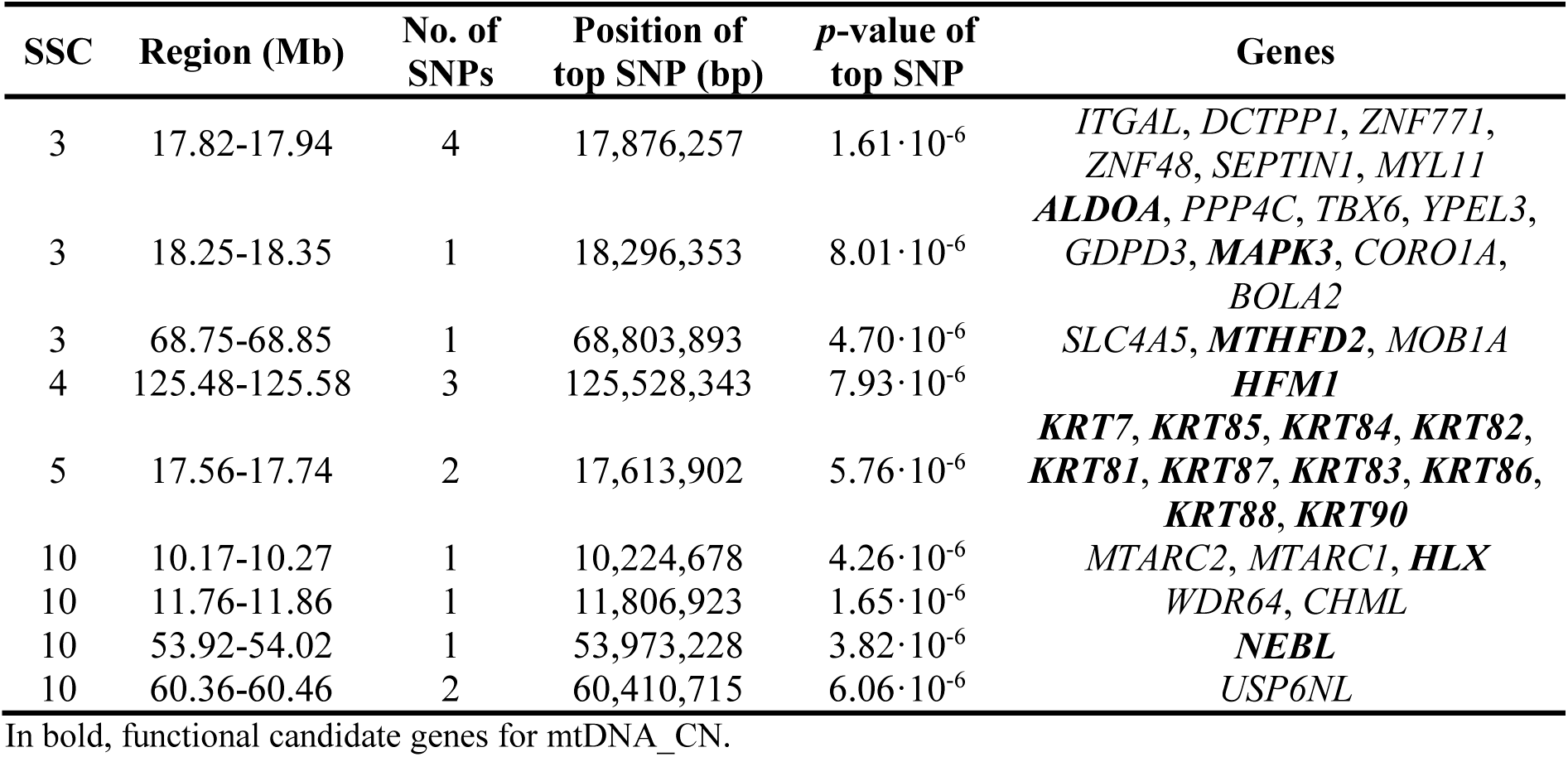
Genomic regions associated with mtDNA_CN in the Duroc population.

## Discussion

We used whole-genome sequence data to determine mtDNA_CN as a proxy of mitochondrial activity in skeletal muscle of Duroc pigs. Our findings suggested that this cellular phenotype exhibited an association with carcase lean percentage. Pigs with lower mtDNA_CN displayed reduced fat content and increased loin mass. Additionally, we analysed the genetic variation within the mtDNA and conducted a GWAS to pinpoint specific genomic regions harbouring promising functional genes that could regulate mitochondrial activity.

### Association between mtDNA_CN and production traits

Our study represents the first attempt to explore the relationship between mtDNA_CN and production traits in pigs. Two previous studies delved into this association in chickens and cattle. In chickens, Reverter *et al*. (2016) employed quantitative polymerase chain reaction (qPCR) to quantify various mtDNA segments, normalizing their abundance against a nuclear DNA segment. Their research spanned different tissues and showed a positive correlation in mtDNA_CN among various muscles, suggesting a coordinated regulation of mitochondrial content across skeletal muscles. Moreover, they found that higher muscular mitochondrial content was linked to lower levels of abdominal fat, breast meat yield, and carcase yield. Another study, conducted by Sanglard *et al*. (2023) in cattle, utilized low-pass sequencing data from blood samples. Their investigation revealed a negative correlation between mtDNA_CN and birth and weaning weights, but not perceptible in mature animals. Both of these studies appear to indicate that reduced mitochondrial activity contributes to an increase in body weight and fat deposition, although, for cattle, this effect was only observable before weaning. These findings align with existing fat metabolism models, which have noted a higher susceptibility to obesity in individuals with lower rates of oxidation (Tanner *et al*., 2002) or low mitochondrial activity (Holmström *et al*., 2012). Moreover, the study of Kaaman *et al*. (2007) concluded that humans with less mtDNA_CN tend to have a higher body mass index (i.e., greater body fat content). In our study, we found that a reduction of mtDNA_CN produced an increase in muscle yield, which is consistent with observations on breast meat yield in poultry. However, contrary to what has been described, we observed a trend towards increased backfat thickness in the pigs with high mtDNA_CN.

In pigs, high feed efficiency lines tend to show less backfat thickness and more loin thickness (Hoque *et al*., 2008; Faure *et al*., 2013). Consequently, our research outcomes support the hypothesis that a reduction of mtDNA_CN or activity is associated with an increase of feed efficiency, producing leaner animals. Although we do not have data on feed intake or feed efficiency, we have demonstrated that a missense variant in the *LEPR* gene affects feed efficiency in our population through interfering with the satiety of the pig (Solé *et al*., 2021). However, we found no relationship between the genotypes for this variant and mtDNA_CN and adding the LEPR genotype into the model did not alter the estimated differences between mtDNA_CN groups (Table S2), indicating independent modes of action.

We found a positive correlation between mtDNA_CN and meat ultimate pH. It is well-established that mitochondria play a role in muscle-to-meat conversion (Hudson, 2012; Popp *et al*., 2015; Dang *et al*., 2020, 2022). Mitochondrial content has been previously associated with dark-cutting beef. Ramanathan *et al*. (2020) observed a greater mitochondrial content and a downregulation of metabolites involved in glycolytic pathways in dark-cutting beef, which can explain the lower lactic acid formation during anaerobic metabolism; thus, resulting in a higher pH. An in vitro study using mitochondria isolated from porcine *longissius lumborum*, detected that mitochondria have a positive impact on ATP maintenance, while mitochondrial inhibitors promote ATP loss, glycogen degradation and lactate accumulation, ultimately reducing the pH (Scheffler *et al*., 2015). The positive correlation we observed aligns with these studies, suggesting that a higher mtDNA_CN is associated with an elevated pH. However, we did not observe any substantial difference for meat ultimate pH between the extreme mtDNA_CN groups.

The cellular phenotype mtDNA_CN is sensible to the sampled tissue from which DNA was extracted. As expected, we detected great differences for mtDNA_CN between skeletal muscle and blood samples (Wachsmuth *et al*., 2016; St. John, 2019; Picard, 2021). However, a similar correlation structure of mtDNA_CN in both tissues with the main production traits suggests that non-target tissues might be similarly used, for example blood (Sanglard *et al*., 2023) or non-target skeletal muscles (Reverter *et al*., 2016). In our study, blood samples were collected at a much earlier age than skeletal muscle samples, which can impact those correlations, which in our case was most noticeable in carcase weight. Thus, the use of non-target tissues for mtDNA_CN estimation should be properly validated before it can be used as a predictor for other performance traits.

### mtDNA_CN genetic background

Mitochondrial genome polymorphisms had been previously related to production traits (Fernández *et al*., 2008; St. John and Tsai, 2018; Liu *et al*., 2023). However, we did not observe any discernible impact of mitochondrial haploblocks on mtDNA_CN or on production traits. This lack of correlation may be attributed to the utilization of a purebred Duroc line in our research, as opposed to commercial F1 or F2 hybrids. Consequently, lower levels of variation in the mtDNA genome within our population might account for the absence of a notable association with production traits.

Nuclear DNA also contributes to variation in mitochondrial activity and mtDNA_CN. We estimated a heritability for mtDNA_CN of 0.60 ± 0.46 using a sire model (Falconer and Mackay, 1996). Despite the large standard error, this estimate falls within the range reported in the literature (Xing et al., 2008; López et al., 2012) and confirms a genetic component for the variance of mtDNA_CN.

The GWAS identified 27 nuclear DNA variants associated with mtDNA_CN. Of those, 16 were located at genomic regions with annotated genes. A literature search of the variants located at those regions revealed several functional candidate genes. We detected two genomic regions in chromosome SSC3. The first one, located at around 18 Mb, contains two genes that had been previously reported to be associated with mitochondria regulation. The first one is the gene coding for aldolase A (*ALDOA*), described as a regulator of mitochondrial respiration in cancerous pancreatic cells (Ji *et al*., 2016). Moreover, it has been recently demonstrated that *ALDOA* restricts PRKN-dependent mitophagy (Bai *et al*., 2022). Mitogen-activated protein kinase 3 (*MAPK3*), also known as extracellular signal-regulated kinase 1 (*ERK1*), is also associated with mitochondrial function; specifically with mitochondrial fission leading to a decrease in mitochondrial mass during cell reprograming (Prieto *et al*., 2016). Other studies have reported that activation of the ERK1/2 signalling pathway produces an inhibition of glycolysis and oxidative phosphorylation, which leads to mitochondrial dysfunction (Nowak, 2002; Zhu *et al*., 2012; Zhou *et al*., 2023). Another region located at 69 Mb of SSC3 has been associated with mtDNA_CN. Among the genes located at that region, methenyltetrahydrofolate cyclohydrolase (*MTHFD2*) stands out. In mouse, a knock-down of *MTHFD2* produces a glucose metabolism shift, from oxidative phosphorylation to lycolysis. This change in the metabolism has been linked to the interaction between this gene and complex III member of the mitochondrial electron transport chain. A depletion of *MTHFD2* reduces considerably the stability of the complex, leading to a mitochondrial disfunction (Yue *et al*., 2020; Zhao *et al*., 2022).

On chromosome SSC4, at around 125 Mb, a signal has been detected. The helicase for meiosis 1 (*HFM1*) gene maps to this region. Although this gene might seem unrelated with mitochondrial function, it has been observed in *Saccharomyces cerevisiae* yeast that the overexpression of *HFM1* is capable to restore respiratory capacity, cytochrome spectrum and oxygen consumption (Tigano *et al*., 2015).

On chromosome SSC5, at 17 Mb, we located a cluster of keratin genes. Keratin, which forms intermediate filaments found in all epithelial cells, plays a critical role for stress protection and cellular integrity. Cells from individuals affected by pachyonychia congenita, a disease caused by mutations in type II keratins 6a, 6b and 6c, and type I keratins 16 and 17, showed a reduction in mitochondrial-endoplasmic reticulum contact sites. The loss of these contact sites has been linked to a reduction in mitophagy, and, consequently, an accumulation of old and dysfunctional mitochondria (Lehmann, Leube and Schwarz, 2020). Additionally, dysmorphic mitochondria were detected in *Krt5* and *Krt16* knock-out mice (Takahashi, Folmer and Coulombe, 1994; Alvarado and Coulombe, 2014), reinforcing its involvement in mitochondrial function.

On chromosome SSC10, at 10 Mb, there is located the gene HLX, which encodes a transcription factor that in mice is co-activated by *Prdm16* to control brown adipose tissue gene expression and mitochondrial biogenesis (Huang *et al*., 2017), and which also regulates mitochondrial metabolic genes (Piragyte *et al*., 2018). Furthermore, in this chromosome we identified another region at 54 Mb where the nebulin gene (*NEBL*) is located. This gene is an important contributor on the maintenance of myofibrillar integrity during muscle contraction. Also, knock-out mice for this gene displayed abnormal accumulation of mitochondria within myofibers (Bang *et al*., 2006).

### Conclusions

Pigs with less mtDNA_CN produce leaner carcases than pigs with high mtDNA_CN. This result supports the hypothesis that a reduction of mitochondrial activity is associated with greater feed efficiency. Pork from pigs with less mtDNA_CN may also tend to have lower ultimate pH as mitochondria play a role in muscle-to-meat conversion. We found no association of the most frequent mitochondrial haploblocks with mtDNA_CN or the production traits, suggesting that mtDNA_CN may be primarily controlled by nuclear DNA. Several genomic regions that harbour potential candidate genes with functions related to mitochondrial biogenesis and homeostasis were associated to mtDNA_CN. These regions provide new insights into the genetic background of this cellular phenotype but it is still uncertain if such associations translate into noticeable effects on the production traits.

## DATA AVAILABILITY

The data that support the findings of this study are available from the corresponding author upon reasonable request.

## CONFLICT OF INTEREST DISCLOUSURE

The authors declare that the research was conducted in the absence of any commercial or financial relationships that could be construed as a potential conflict of interest.

## ETHICAL APPROVAL

The pigs used in this study were raised and slaughtered in commercial units complying with the regulations and good practice guidelines on the protection of animals kept for farming purposes, during transport and slaughter (Royal Decree 37/2014, Spain). Tissue samples used in this study were collected at the slaughterhouse. The Ethical Committee on Animal Experimentation of the University of Lleida approved all experimental procedures (CEEA 04-06/21).

## ACKNOWLEDGMENTS

We acknowledge the personnel at Selección Batallé for their cooperation for the recording of on-farm data and sample collection. We gratefully acknowledge Pilar Sopeña from the Animal Breeding group, University of Lleida, for laboratory assistance.

## CREDIT AUTHOR STATEMENT

EM: Conceptualization, Methodology, Formal analysis, Investigation, Writing - Original Draft. RNP: Resources, Writing - Review and Editing. JE: Resources, Writing - Review and Editing, Funding Acquisition. RRF: Conceptualization, Methodology, Resources, Writing - Original Draft, Supervision, Funding Acquisition.

## Supplementary Tables

**Table S1.**
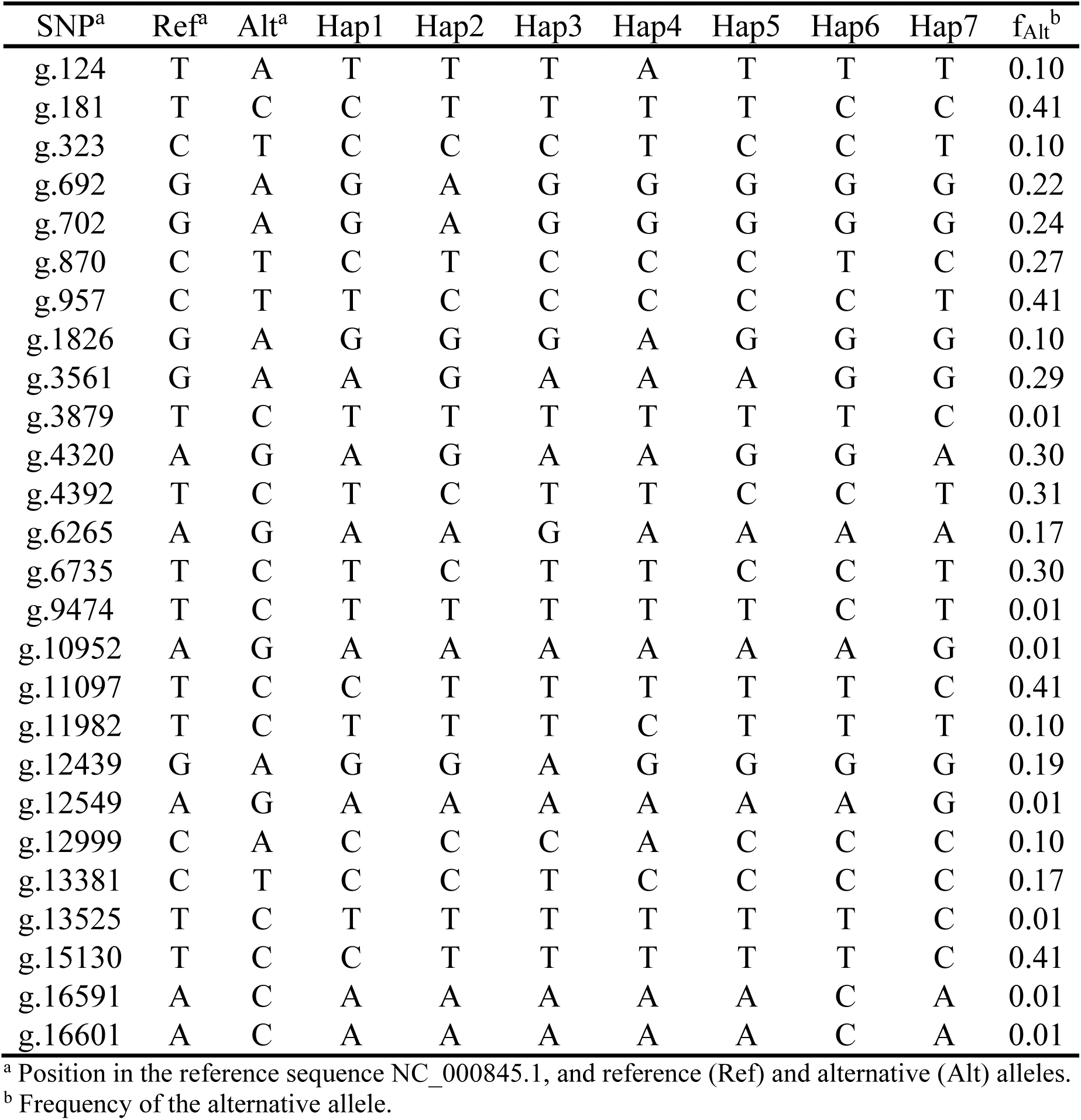
Haplotype frequencies of the single nucleotide polymorphisms (SNP) located at the mitochondrial DNA.

| SNP<sup>a</sup> | Ref<sup>a</sup> | Alt<sup>a</sup> | Hap1 | Hap2 | Hap3 | Hap4 | Hap5 | Hap6 | Hap7 | f<sub>Alt</sub><sup>b</sup> |
| --- | --- | --- | --- | --- | --- | --- | --- | --- | --- | --- |
| g.124 | T | A | T | T | T | A | T | T | T | 0.10 |
| g.181 | T | C | C | T | T | T | T | C | C | 0.41 |
| g.323 | C | T | C | C | C | T | C | C | T | 0.10 |
| g.692 | G | A | G | A | G | G | G | G | G | 0.22 |
| g.702 | G | A | G | A | G | G | G | G | G | 0.24 |
| g.870 | C | T | C | T | C | C | C | T | C | 0.27 |
| g.957 | C | T | T | C | C | C | C | C | T | 0.41 |
| g.1826 | G | A | G | G | G | A | G | G | G | 0.10 |
| g.3561 | G | A | A | G | A | A | A | G | G | 0.29 |
| g.3879 | T | C | T | T | T | T | T | T | C | 0.01 |
| g.4320 | A | G | A | G | A | A | G | G | A | 0.30 |
| g.4392 | T | C | T | C | T | T | C | C | T | 0.31 |
| g.6265 | A | G | A | A | G | A | A | A | A | 0.17 |
| g.6735 | T | C | T | C | T | T | C | C | T | 0.30 |
| g.9474 | T | C | T | T | T | T | T | C | T | 0.01 |
| g.10952 | A | G | A | A | A | A | A | A | G | 0.01 |
| g.11097 | T | C | C | T | T | T | T | T | C | 0.41 |
| g.11982 | T | C | T | T | T | C | T | T | T | 0.10 |
| g.12439 | G | A | G | G | A | G | G | G | G | 0.19 |
| g.12549 | A | G | A | A | A | A | A | A | G | 0.01 |
| g.12999 | C | A | C | C | C | A | C | C | C | 0.10 |
| g.13381 | C | T | C | C | T | C | C | C | C | 0.17 |
| g.13525 | T | C | T | T | T | T | T | T | C | 0.01 |
| g.15130 | T | C | C | T | T | T | T | T | C | 0.41 |
| g.16591 | A | C | A | A | A | A | A | C | A | 0.01 |
| g.16601 | A | C | A | A | A | A | A | C | A | 0.01 |

<sup>a</sup> Position in the reference sequence NC\_000845.1, and reference (Ref) and alternative (Alt) alleles.
<sup>b</sup> Frequency of the alternative allele.

**Table S2.** Least-square means (± standard errors) for production traits by mtDNA_CN group accounting for *LEPR* genotype in the model.

| Trait | Low | High | *p*-value |
| --- | --- | --- | --- |
| Live measurements at 207 d |  |  |  |
| Body weight, kg | 128.4±2.2 | 126.3±2.3 | 0.54 |
| Backfat thickness, mm | 21.4±0.6 | 23.7±0.7 | **0.04** |
| Loin thickness, mm | 48.5±1.0 | 48.2±1.1 | 0.82 |
| Carcase measurements |  |  |  |
| Carcase weight, kg | 98.6±1.4 | 98.2±1.6 | 0.81 |
| Backfat thickness, mm | 22.7±0.5 | 24.0±0.5 | **0.05** |
| Loin thickness, mm | 47.0±1.2 | 42.7±1.3 | **<0.01** |
| Carcase lean percentage, % | 42.8±0.8 | 39.9±0.9 | **0.01** |
| Intramuscular fat content, % | 18.3±0.8 | 19.1±0.9 | 0.53 |
| pH | 5.76±0.04 | 5.80±0.04 | 0.46 |

In bold, *p*-values < 0.10.

